# Genetic differences between *Coccidioides* spp. and closely related nonpathogenic Onygenales

**DOI:** 10.1101/413906

**Authors:** Theo N. Kirkland

## Abstract

*Coccidioides* spp. are dimorphic, pathogenic fungi that can cause severe human and animal disease. Like the other primary fungal pathogens, animal infection results in a morphologic transformation from the environmental mycelial phase to a tissue phase, known as a spherule. The sequencing and annotation of *Coccidioides* spp. and the genomes of several nonpathogenic Onygenales species allows comparisons that provide clues about the *Coccidioides* spp. genes that might be involved in pathogenesis. The analysis in this study is a gene by gene orthology comparison. Although there were few differences in the size of genes families in the *Coccidioides* spp.-specific group compared to the genes shared by *Coccidioides* spp. and nonpathogenic Onygenales, there were a number of differences in the characterization of the two types of genes. Many more *Coccidioides* spp.-specific genes are up-regulated expression in spherules. *Coccidioides* spp.-specific genes more often lacked functional annotation and were more often classified as orphan genes. Analysis by random forest machine learning confirmed that high numbers of orthologs and high levels of expression in hyphae were predictive of common genes, while high levels of expression in spherules and more nonsynonymous predicted *Coccidioides* spp.-specific genes. Review of individual genes in the *Coccidioides* spp.-specific group identified a histidine kinase, two thioredoxin genes, a calmodulin gene and ureidoglycolate hydrolase. Hopefully, identification of these genes will be useful for pursuing potential *Coccidioides* spp. virulence genes in the future.

## Introduction

*Coccidioides immitis* and *Coccidioides posadasii* are dimorphic fungi that are found in the soil in the desert regions of the American southwest, Mexico and South America [1-3]. Both species can cause disease known as Valley Fever in normal people. The two species are closely related, morphologically indistinguishable and the major phenotypic difference between the two that is currently recognized is their geographic distribution; *C. immitis* is found primarily in California (including Baja California) and *C. posadasii in* Arizona and other endemic areas [4,5]. The organism grows as mycelia in the soil and forms arthroconidia within the mycelium that are dispersed as the soil is disturbed by wind, construction or other events. If the arthroconidia are inhaled by susceptible hosts they undergo transformation into a spherical tissue form, known as a spherule. These structures divide internally to form as many as 100 endospores, which in turn can differentiate into spherules. Endospores can be released 96 hours after initial spherule formation, so the number of spherules can increase quickly. Infection with *Coccidioides* spp. causes symptomatic disease in 40-50% of immunocompetent hosts and in a substantial number of infections can be prolonged and/or severe. In some instances, dissemination of the infection beyond the lung occurs. Some infections require life-long treatment and severe infections can be fatal.

*Coccidioides* spp. are a member of the Ascomycete Onygenales order which contains a number of other human pathogenic fungi, including *Histoplasma capsulatum, Blastomyces dermatitidis, Paracoccidioides* spp., and the dermatophytes such as *Microsporum* spp. and *Trichopyton* spp.[6]. *Coccidioides* spp. are in the Coccidioides family, in contrast to *Histoplasma capsulatum, Blastomyces* spp., and *Paracoccidioides* spp. which are all Ajellomycetacae [7]. In addition to these pathogens, many fungal species that are not human pathogens are also Onygenales [6,8]. One of these, *Uncinocarpus reesii*, forms arthroconidia within mycelia that are morphologically indistinguishable from *Coccidioides* spp. [9]. *U. reesii* is found in the soil and has a worldwide distribution. It is keratinophilic and thermotolerant [10,11]. *U. reesii* utilizes amino acids and peptides as well as carbohydrates for growth [12]. *U. reesii* does not cause human disease but a close relative, *Chrysosporium zonatum*, has been reported to cause an infection in a single patient with chronic granulomatous disease [11]. Despite the difference in lifestyle, *U. reesii* is closely genetically related to *Coccidioides* spp. [8,9,13].

The genome sequencing of multiple isolates of *Coccidioides* spp. and one isolate of *U. reesii* has made it possible to identify genes that are present in *Coccidioides* spp. but not this close nonpathogenic relative. At least four other nonpathogenic species in the Onygenales are also closely related to *Coccidioides* spp. [6]. The genomes of these organisms, *Amauroascus mutatus, Amauroascus niger, Byssoonygena creatinophilia, Chrysosporium queenslandicum* and *Onygena corvina* have also been sequenced and the size of gene families and the gain or loss of genes compared to *Coccidioides* spp. and *U. reesii* and other fungi [6]. Whiston and Taylor found about 800 *Coccidioides* spp. genes that were not present in the nonpathogenic Onygenales. One distinctive feature of the *Coccidioides* spp.-specific genes was the increased frequency of preferential expression in spherules [6].

Comparison of the genes conserved within *Coccidioides* spp. but not found in closely related nonpathogenic Onygenales to those shared between *Coccidioides* spp. and nonpathogenic Onygenales might provide insights into genes that are required for pathogenesis of coccidioidomycosis. The purpose of the study is to identify these genes and analyze their characteristics.

## Methods

### Orthology

Initial ortholology searches were done using the orthology tool at FungiDB (http://fungidb.org/fungidb/showApplication.do). This tool compare two sets of proteins by BLASTP and computes the percent match length. The thresholds for blast results were an e value < 10^−5^ and a percent match length of ≥ 50%. Paralogs, orthologs and ortholog groups were found using OrthoMCL Pairs. Orthologs shared by *C. immitis* RS, H538.4 and *C. posadasii* Silvera and RMSCC 3488 were identified. Sequences for all those strains are available at FungiDB. The sequence of *C. immitis* WA_211 is available at NCBI (https://www.ncbi.nlm.nih.gov/nuccore/RHJW00000000). Orthologs of *Coccidioides* spp. genes to *C. immitis* WA_211were identified using the online toll OrthoVenn2 (https://orthovenn2.bioinfotoolkits.net/home). The 7627 genes conserved within these *C. immitis* and *C. posadasii* isolates were then compared to *U. reesii* using the FungiDB orthology tool and reciprocal BLAST with a cut off e value of 10^−6^. The reciprocal blast tool was Reciprocal-BB, which is available from https://github.com/marcusgj13/Reciprocal-BB. Comparison to the predicted proteins of non-pathogenic Onygenales, *A. mutatus, A. niger, B. creatinophilia, C. queenslandicum* and *O. corvina* was also done with Reciprocal-BB, with a cut off e value of 10^−6^. The gene predictions and annotations for these species were provided by Jason Stajich, using DNA sequence data from Whiston and Taylor [14]. The *C. immitis* R.S. gene I.D. notation is used for all gene designations.

### Blast

BLASTP searching was done using Stand-alone BLAST from NCBI, using the default settings. Blast hits were identified using an upper limit e value of 10^−6^. Searches against fungal proteins were done using a representative sample (3 million predicted proteins) of the fungal protein database downloaded from NCBI. *Coccidioides* spp. genes without BLASTP matches in this search were defined as orphan genes.

### Gene characterization

Other characteristics, such as the genetic location, ortholog group, number of orthologs, predicted signal peptides, predicted transmembrane domains, PFAM and other functional domains. The functional domains included data from UniProt (www.uniprot.org), Superfamily 1.75 (http://supfam.org/), TIGR(https://www.jcvi.org/institute-genomic-research-tigr-j-craig-venter-institute-j-craig-venter-science-foundation) and NCBI (https://www.ncbi.nlm.nih.gov/cdd). The relative levels of expression in spherules and mycelia were downloaded from FungiDB. Up-regulation was defined by the criteria described by Whiston et. al., comparing mycelia to day 4 spherules [14,15]. The Chi-squared test was used to determine the statistical significance of differences between ratios; the Kruskal-Wallis test was used to compare determine the statistical significance of differences between continuous variables. All data analysis was done in R.

## Supplementary data

Supplemental Table 1 is an attached Excel table containing all the data used for these analyses.

## Results

7627 *Coccidioides* spp. genes are shared by *C. immitis* RS and H538.4, WA_211 and *C. posadasii* Silvera and RMSCC 3488. Since two strains of *C. immitis* from southern California and a *C. immitis* strain from Washington [16], as well two strains of *C. posadasii* were used to select this group, most of the core genome of *Coccidioides spp*. genome should be included. However, many more strains have been sequence but not annotated, so this may be an over-estimate of the total number of conserved genes [17]. The total number of genes in these isolates varies between 9,905 and 117,815 (mean = 9,768) so, on average, about 20% of the genes appear to be strain specific. They will not be considered further here.

632 conserved *Coccidioides spp*. genes are not found in *U. reesii* or *A. mutatus, A. niger, B. creatinophilia, C. queenslandicum* and *O. corvina*. This value was similar to the estimate previously obtained by Whiston and Taylor [14]. For brevity, these genes are referred to as *Coccidioides* spp.-specific genes and those that are common to the *Coccidioides* spp. and nonpathogenic Onygenales are referred to as common genes. The term *Coccidioides* spp.-specific does not imply that these genes do not have homologs in other fungi.

Table 1 shows a comparison of the two sets of genes. Ninety-two percent of all the *Coccidioides* spp. genes are common to nonpathogenic Onygenales and 8 % are *Coccidioides* spp.-specific genes. The predicted properties of proteins in the two groups are similar except for homology to other fungal proteins, the preferential expression in spherules, the amount of functional annotation and the degree of genetic variation. The mean number of orthologs per gene, the percentage of genes with blast matches, PFAM or other functional domains or functional product descriptions (not annotated as “hypothetical” or “predicted protein”) are all higher in the common genes than the *Coccidioides* spp.-specific genes. As would be expected, common genes are homologous to genes in a wider diversity of fungi. There was no significant differences between the two groups in the proportion of genes with GO terms but only a small percentage of either group had a predicted GO term. Although many genes in both groups had a PFAM domain or other functional domain, very few terms were present in significant numbers. A larger proportion of the *Coccidioide*s spp.-specific genes are expressed at a higher level in spherules than mycelia, indicating that this group of genes is enriched for genes associated with differentiation to spherules. 30% of the *Coccidioides* spp.– specific genes have no BLASTP matches to other fungi with an e value less that 10^−6^, which is our definition of orphan genes. 45% of the orphan genes are up-regulated in spherules. The origin of orphan genes is not clear. Although the function of these genes is difficult to determine, their conservation within *Coccidioides* spp. and up-regulation in spherules suggests that they may be important in differentiation into spherules. The *Coccidioides* spp.-specific genes did not cluster on the genetic map.

**Table 1.** Comparison of *Coccidioides* spp.- specific to common genes. a: Compared to total number of *Coccidioides* spp. genes conserved within the *C. immitis* and *C. posadasii* strains tested; b: Compared to total number of *Coccidioides* spp.- specific genes; c: based on annotation of predicted proteins; d: Nonsynonymous/synonymous single nucleotide polymorphisms.

| Characteristics | Number (%) |  |  |  | P |
| --- | --- | --- | --- | --- | --- |
|  | *Coccidioides* spp.- specific |  | Common |  |  |
| Total | 632 | 8<sup>a</sup> | 6955 | 92<sup>a</sup> | <10<sup>-10</sup> <sup>b</sup> |
| Mean orthologous groups/gene | 47 |  | 107 |  | <10<sup>-10</sup> |
| Up-regulated in spherules | 184 | 29 | 911 | 13 | <10<sup>-10</sup> |
| Expression unchanged | 363 | 57 | 4985 | 72 | <10<sup>-10</sup> |
| Down-regulated in spherules | 85 | 13 | 1059 | 15 | N.S. |
| Orphan genes | 190 | 30<sup>b</sup> | 0 | 0 | <10<sup>-10</sup> |
| Functional product description <sup>c</sup> | 211 | 34 | 4624 | 66 | <10<sup>-10</sup> |
| Functional domains | 331 | 52 | 5750 | 83 | <10<sup>-10</sup> |
| Mean NS/S<sup>d</sup> | 15.2 |  | 2 |  | <10<sup>-10</sup> |
| One or more transmembrane region | 104 | 16 | 1313 | 19 | N.S. |
| Signal peptide | 78 | 45 | 1594 | 25 | N.S. |

A previous study has identified 26 genes that are up-regulated in *Coccidioides* spp. and *Histoplasma capsulatum* as the spherule or yeast differentiation occurs [18]. Four of these are also found in the up-regulated *Coccidioides* spp.-unique group. One of the most interesting of these is CIMG_02628, an ARP2/3 complex subunit. This complex is composed of seven subunits that control nucleation of actin, which is important for cell wall remodeling and endocytosis [19]. The transformation of mycelia to spherules requires extensive cellular remodeling, so the up-regulation of this gene seems reasonable. Two GMC/SPRK kinases are up-regulated *Coccidioides* spp.-specific genes; one of these, CIMG_02373, is also up-regulated in *H. capsulatum* yeast.

### Analysis of individual *Coccidioides* spp.-specific genes

Because the number of *Coccidioides* spp.-specific genes was relatively small, individual genes of interest could be found by inspection. A number of these are shown in Table 2. Many other genes may also be of interest but these seem the most promising for further study.

**Table 2.**
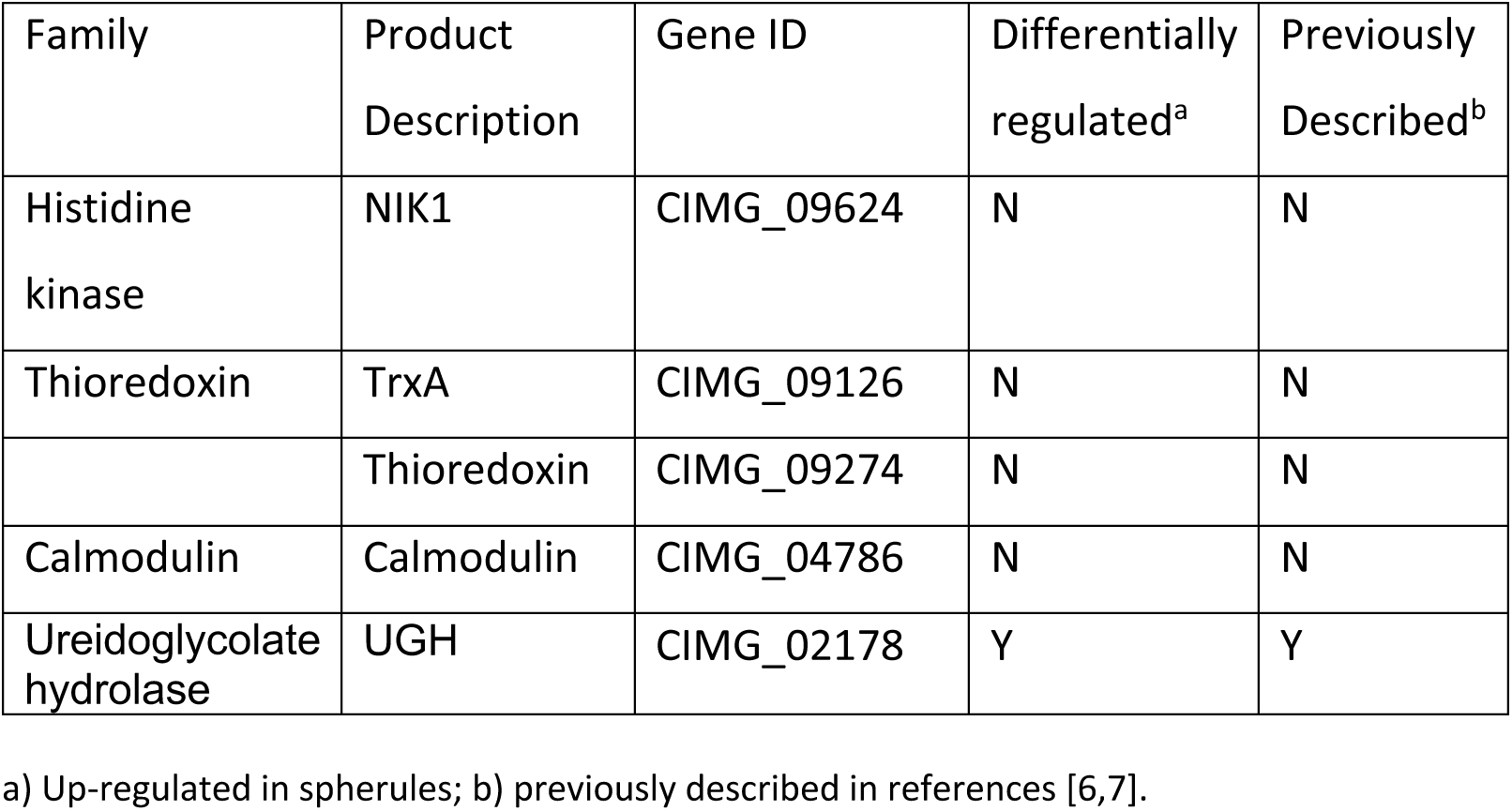
*Coccidioides* spp.-specific genes of interest

### Histidine kinase

Histidine kinases are another group of proteins involved in responses to external stimuli [20]. Some of the best know functions of histidine kinases include osmosensing, adaption to oxidants, conidiation, cell wall integrity and virulence. In some other primary pathogenic fungi, the Drk1 has been found to be required for the temperature dependent conversion from mycelia to yeast [21,22]. The Drk1 gene is found in both *Coccidioides* spp. and nonpathogenic Onygenales.

Goldberg described three histidine kinases in *C. immitis*, CIMG_04512, CIMG_05052 and CIMG_06539, which are all present in *U. reesii* [15]. However, there are more than ten histidine kinases in most filamentous fungi, so it is likely that more genes exist [23]. In this analysis, we identified one other *Coccidioides* spp.-specific gene with a histidine kinase domain, which is not up-regulated in spherules. This gene, CIMG_09624, is almost completely identical in many isolates of *Coccidioides* spp. and very similar to histidine kinases in several other taxa despite its absence in non-pathogenic Onygenales. These data argue that this gene is very probably a conserved histidine kinase that may play an important role in the biology of *Coccidioides* spp.

### Thioredoxin

Thioredoxin is an important regulatory enzyme that modifies target proteins by modifying their redox status [24]. Most organisms have multiple thioredoxin genes with overlapping functions. The biological role of thioredoxin includes antioxidant functions that are probably particularly important for *Coccidioides* spp. when ingested by phagocytic cells. *C. immitis* is relatively resistant to oxidative stress and mice lacking an oxidative burst are not more susceptible than control mice to coccidioidomycosis [25]. There are a total of three thioredoxin genes in *Coccidioides* spp., CIMG_09126, CIMG_09274 and CIMG_08211, none of which are up-regulated in spherules. Two of these, CIMG_09126 and CIMG_09274, are not found in nonpathogenic Onygenales but *U. reesii* does has have an ortholog of CIMG_08211. This difference in the number of thioredoxin genes may play a part in the ability of *Coccidioides* spp. to survive an oxidative burst and survive phagocytosis.

### Calmodulin

Calmodulin is a calcium binding protein in the calcineurin pathway which is also important in regulating growth, responses to stress and virulence in a variety of fungi [26,27]. When calmodulin binds to calcium it activates calcineurin, which is critical for thermotolerance and pathogenicity. Calcineurin inhibitors inhibit the growth of a number of different fungi, including *C. immitis*, at 37°C [27,28]. There are three genes coding for calmodulin in *Coccidioides* spp. and one of these, CIMG_04786 is a *Coccidioides* spp.-specific gene. *U. reesii* has only two genes coding for calmodulin. This difference in the number of genes may play a role in determining thermotolerance.

### Ureidoglycolate hydrolase

Ureidoglycolate hydrolase (*UGH*), CIMG_02178, is a *Coccidioides spp*.-specific gene. Cole and his co-workers have studied the role of extracellular ammonia in spherule development [29]. There is evidence that first generation spherules release endospores into an alkaline environment [29]. One enzyme involved in ammonia production is *UGH* and the expression of this gene is up-regulated in spherules compared to mycelia [14]. The *C. posadasii* UGH deletion mutant is less virulent than the wildtype, even though the mutant grew normally as a mold in vitro [29]. The urease/UGH double mutant, which makes very little ammonia, was much less virulent than wildtype. These data suggest that the UGH gene plays an important role in the pathogenesis of infections caused by *Coccidioides* spp.

### Comparison to other studies

As part of a study of the genome of pathogenic Ajellomyces fungi compared to close non-pathogenic relatives Cuomo identified 46 *C. immitis* genes that were not found in *U. reesii* [7]. The current analysis identified many more genes that were found in multiple *Coccidioides* spp. isolates but not nonpathogenic Onygenales but 24 of the genes were found in both studies. Cuomo also proposed 32 genes that were important for fungal pathogenesis. This analysis found only two homologs of these genes in Coccidioides spp. specific group: Ureidoglycolate hydrolase (*UGH*) and Ryp2. Whiston and Taylor have also compared the predicted proteins of *Coccidioides* spp. to nonpathogenic Onygenales [6]. The number of *Coccidioid*es spp.-specific genes and the tendency of those genes to be over-expressed in spherules is similar in their study and this one.

### Summary

This attempt to identify *Coccidioid*es spp. genes that are not found in nonpathogenic Onygenales is done to try to identify candidate pathogenesis genes. The strengths of the study include the overlap of these results with previous studies and the internal consistency of the data. This study used slightly different methods than others but a significant number of *Coccidioides* spp.-specific genes were found in more than study. There are also a number of weaknesses. The identification of unique and common genes depends on the accuracy of the current annotation. Since *U. reesii* and the other nonpathogenic Onygenales may not be annotated as well as *C. immitis* and *C. posadasii*, some genes might be classified as unique in error. Furthermore, a weakness of all these types of studies is the possibility that genes have evolved to perform new functions, so the absence of a given homolog in an organism does not imply the absence of that biological process. In addition, in the instances where a number of genes are found (such as thioredoxin and calmodulin) it is not clear what to expect from the result of the absence of one or two of them.

Many of the unique genes are not only divergent from non-pathogenic Onygenales, but lack homologs in other fungal taxa. 30% of the Coccidioides spp.-specific genes are orphans but none of common genes are. These orphan genes are found in at least five isolates of *Coccidioides* spp. and many are up-regulated in spherules.

A common theme of the genes that are *Coccidioides* spp.-specific is their role in responding to environmental changes. In addition, *Coccidioides* spp.-specific genes tended to be up-regulated in spherules more often than *Coccidioides* spp. shared with nonpathogenic Onygenales, suggesting that more *Coccidioides* spp.-specific genes are involved in, or a consequence of, spherule development. Because many of the up-regulated *Coccidioides* spp.-specific genes are orphan genes their role in mycelium to spherule transformation is intriguing even though their function is unknown. Hopefully this data will suggest genes that would be good candidates for further investigation of the molecular mechanisms of *Coccidioides* spp. pathogenicity.

## Acknowledgement

The assistance of Jason Stajich in planning this study, providing the unpublished annotations and interpreting the results is deeply appreciated.

